# Elevated gene expression of most microglial markers, and reduced expression of most pyramidal neuron and interneuron markers, in postmortem autism cortex

**DOI:** 10.1101/032557

**Authors:** Rebeca Borges-Monroy, Chris P. Ponting, T. Grant Belgard

## Abstract

Autism Spectrum Disorders (ASD) are clinically and genetically heterogeneous. Nevertheless, a characteristic gene expression signature in postmortem cortex is shared across most ASD cases. Knowing whether this signature reflects changes in cells of particular types would help to determine the molecular, cellular, and anatomical etiology of ASD. To investigate we took advantage of cell type marker genes defined by recent single cell transcriptome sequencing from mouse or human cortex. We find that microglial markers showed significantly and substantially higher median expression in postmortem ASD than in control cortex for both mouse and human markers (21% higher for orthologous markers defined in mouse; 93% higher for human markers). In contrast, neuronal markers showed reduced median expression: 12% and 16% lower for mouse interneurons and pyramidal neurons, 40% lower for human neurons, and 36% lower for a class of human excitatory projection neurons. Cell type density alterations in ASD brains are indicated because distributions of these cell type markers are shifted concordantly. Importantly, orthologous mouse neuronal marker genes encoding proteins localized to diverse neuronal compartments all showed reduced median expression in ASD. Our results provide a framework for revealing the basis of transcriptomic differences in ASD. We propose comparing cell type density alterations in post-mortem tissue with and without these distinctive gene expression changes to reconcile results from stereological and transcriptomic studies.

## Introduction

Autism Spectrum Disorders (ASD) are heterogeneous and highly heritable neurodevelopmental disorders characterized by verbal and nonverbal communication deficits, impaired social interactions, and repetitive behaviors from early childhood^1–3^. Despite its strong heritability, genetic susceptibility to ASD in the population is distributed over a thousand genes^4^, which complicates efforts to generalize from deep functional studies of rare but highly penetrant variants. Given this panoply of genetic causes, and a wide variety of proposed causal mechanisms for ASD, it was unexpected that most post-mortem brains from people with ASD exhibit similar gene expression profiles^5^. By studying this characteristic gene expression signature we seek to reveal neurological features that are common to much of the ASD spectrum.

We are only beginning to understand this molecular signature. Alternative splicing and the attenuation of regional gene expression differences in cortex appear to be key components^5, 6^. Nevertheless, there are also early signs that larger differences exist with regard to the density of certain cell types or subcellular components. In particular, a set of co-expressed genes underexpressed in ASD is enriched in synaptic genes^5^, ASD risk genes^5, 7^ and markers of glutamatergic neurons^5, 7^, and contains some interneuron markers^5^. A set of co-expressed genes overexpressed in ASD is enriched for immune-related genes, markers for activated microglia, and markers for astrocytes^5^. Another study found an overrepresentation of genes overexpressed in ASD in immune cell types and astrocytes and an overrepresentation of genes underexpressed in ASD in neuronal cell types, particularly in interneurons^8^.

A parsimonious neurological interpretation of cell type marker enrichment, nevertheless, remains unclear. Such enrichments might be driven by a subset of markers of that type and could therefore reflect changes in cell signaling, physiological state, functional pathways, structure, and more. For example, defects in synapse formation in ASD^9, 10^ may underlie the underexpression of a subset of neuronal marker genes expressed at the synapse. On the other hand, if most neuronal markers are underexpressed in ASD, this could also suggest a reduction in neuronal density. Here we use density to refer to cell number density (the number of cells in a volume of tissue)^11^ or other ratios such as volume density (the total volume of cells in a volume of tissue)^12, 13^ or cross-sectional area stained with a marker (e.g. Nissl-stained area to unstained area in a section)^14^. Indeed, there are scattered and sometimes conflicting reports of changes in neuronal density, which may reflect small sample size and the heterogeneity of ASD^14–19^. Identifying the subset of post-mortem brains in which these changes are expected to occur and the cell types involved in these alterations could guide future confirmatory stereological studies in a more targeted manner.

Previous studies have used a biased subset of cell type marker genes from a biased subset of cell types isolated with pre-selected markers followed by bulk gene expression profiling^8, 20^. However, clustering cells by type without prior information about their markers is now feasible with single cell transcriptomics^21^. Objective, comprehensive, and selective cell type markers can be defined from these clusters^22^. In this study, we examine how orthologs of mouse cell type markers^23^ are shifted in their expression in postmortem ASD brains. Furthermore, we obtain similar findings with human cell type markers from a second single cell transcriptome study^6^.

## Materials and Methods

### Datasets and Gene Lists

#### Expression Data

We used normalized, regressed, log_2_ transformed post-mortem gene expression values from the superior temporal gyrus of 12 ASD cases and 12 matched controls, ranging from 15 to 60 years old and of both sexes^6^. The cases and controls in that study were 57% of the individuals chosen from a larger unpublished RNA sequencing cohort on the basis that they display the strongest of this characteristic gene expression signature.

#### Cell Type Marker Genes

Mouse cell type marker genes were taken from Table S1 of Zeisel et al.^23^. They were based on RNA sequencing of single cells isolated on a Fluidigm C1 AutoPrep System from the somatosensory cortex and CA1 of hippocampus from mice of both sexes between postnatal days 21 and 31^23^. Human cell type marker genes, as well as marker genes of neuronal communities, were taken from Table S3 of Darmanis et al.^24^ and were based on RNA sequencing of single cells isolated on a Fluidigm C1 from freshly resected temporal cortex of several humans of both sexes aged 21 through 63 years^24^. The numbers of these marker genes are provided in Supplementary Table 1.

#### Localization of Neuronal Marker Genes

We used S1 pyramidal marker genes from a single cell transcriptomic study from mouse brain tissue^23^ having 1:1 mouse to human orthologs as defined in Ensembl. The human Ensembl identifiers for these transcripts gave linked subcellular protein localization information for their encoded proteins from the UniProt database^25^. Markers were separated into groups based on whether they contained the following Gene Ontology Cellular Component identifiers: "GO:0005634" for nuclear protein localization, "GO:0030425" for protein localization to dendrites, and "GO:0030424" for protein localization to axons^26^. Most transcripts do not have only one localization identifier. Since there was a considerable overlap between transcripts with the axonal localization identifier and the dendritic localization identifier, these were analyzed together as a single group. To improve specificity, transcripts with both nuclear localization identifiers and axonal or dendritic identifiers were removed from the analysis. We analyzed the median log_2_(expression fold change) of these different subsets of mouse orthologous S1 pyramidal neuron and interneuron marker genes as described below. Caveats of this method include that it assumes that a change in density of neuronal processes (and only processes) would be reflected in the expression levels of transcripts whose proteins are localized to neuronal processes but would not be reflected to a comparable extent in the expression levels of transcripts whose proteins are localized to the cell soma^27, 28^, that there is conservation of protein localization for these markers from mouse to human, and that the ontology’s cellular localization evidence may require further validation^26^.

### Examining Cell Type Gene Expression Changes in ASD

For the mouse cell type markers, we cross-referenced the mouse gene symbols to Ensembl Mouse Gene identifiers, and then took the 1:1 mouse-human orthologs that were also included in the ASD sequencing study^6^ using Ensembl BioMart^29^. For the human cell type markers, we kept those markers that were in the ASD sequencing study. We plotted the distributions of their log_2_ fold changes with R. We tested the two-tailed significance of the median log_2_ fold change for each cell type by constructing a null distribution by permuting case-control labels 1000 times and then subtracting the mean log_2_ (FPKM) in controls from the mean log_2_ (FPKM) in cases. These were corrected for multiple testing using the “p.adjust” function in R^30^, applying the Benjamini and Hochberg False Discovery Rate (FDR) method^31^ with an FDR cut-off of 5%. Confidence intervals were calculated from 1000 bootstraps.

## Results

We first tested a null hypothesis that the median expression fold change for each of eight high-level cortical cell types defined in mouse^23^ was not significantly different from the change observed when case-control labels are randomly shuffled Figure 1). To do so we considered first a set of marker genes defined by single cell sequencing in mouse cortex, before then obtaining similar results for marker genes of six high-level cell types defined in human cortex (Figure 2; Supplementary Tables 2–3). Compared to controls, the median expression of mouse-defined gene markers for S1 pyramidal neurons was significantly lower (by 10–21% at an FDR<5%^31^) in ASD post-mortem human cortex tissue; similarly, median expression of mouse-defined interneuron markers was reduced in ASD brains by 8–16%. By contrast, median expression of mouse-defined microglia markers was increased in ASD brains by 15–28%. Ependymal, endothelial, and mural marker genes also exhibited slightly increased expression ( at a slightly relaxed False Discovery Rate^31^ of 5.1%).

**Figure 1.**
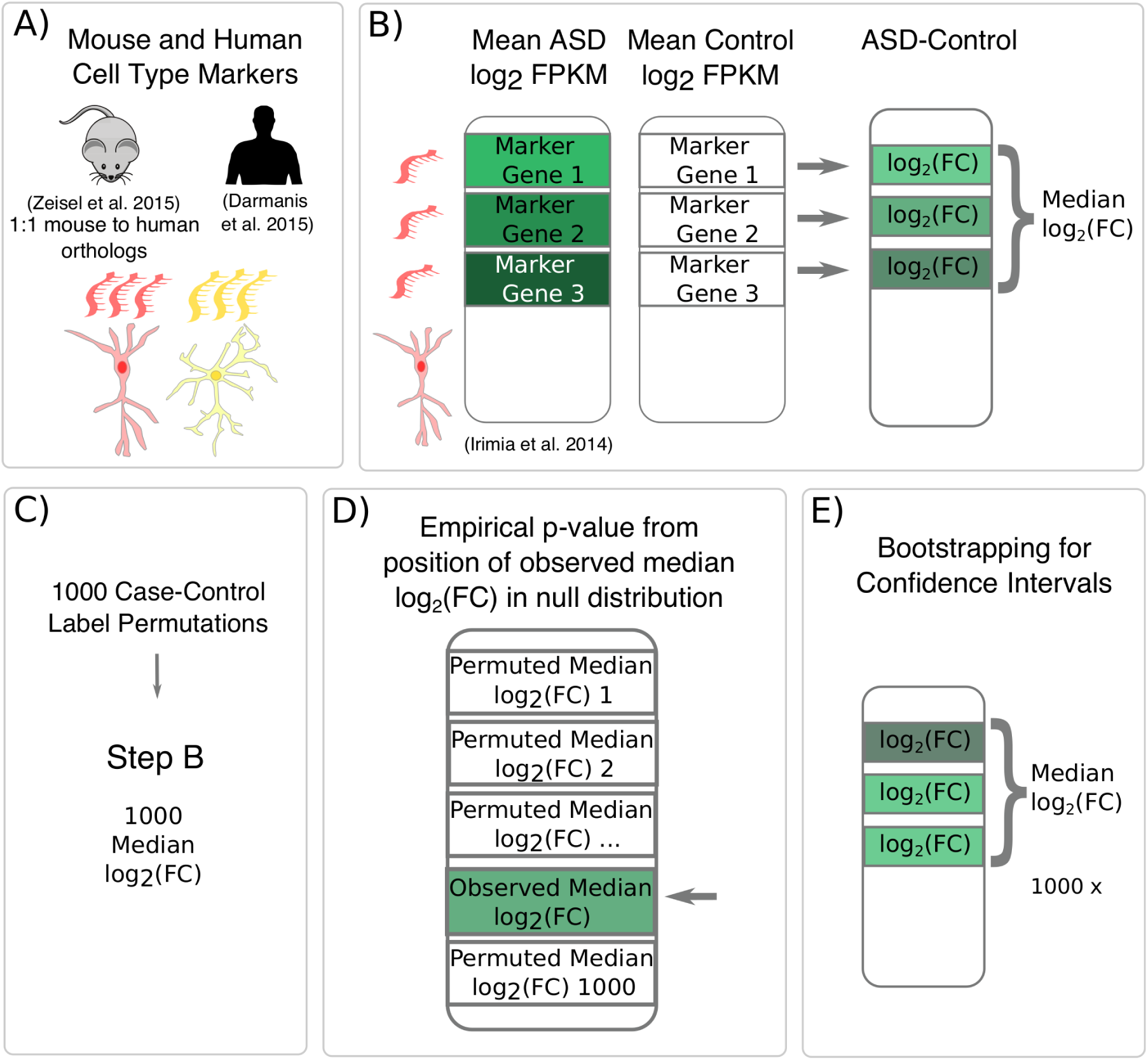
Schematic describing the analyses. A) Cell type markers were obtained from mouse^23^ and human^24^ single cell sequencing studies. B) The median expression log_2_(Fold Change [FC]) in ASD was calculated for all marker genes in each cell type. Genes underexpressed in ASD are indicated in green. C) Case-control labels were permuted 1000 times to compute a null distribution of median log_2_(FC). D) An empirical p-value for the observed median log_2_(FC) was obtained from its position in this null distribution. E) 95% Confidence intervals were obtained by bootstrapping log_2_(FCs) of each cell type and obtaining a median 1000 times.

**Figure 2.**
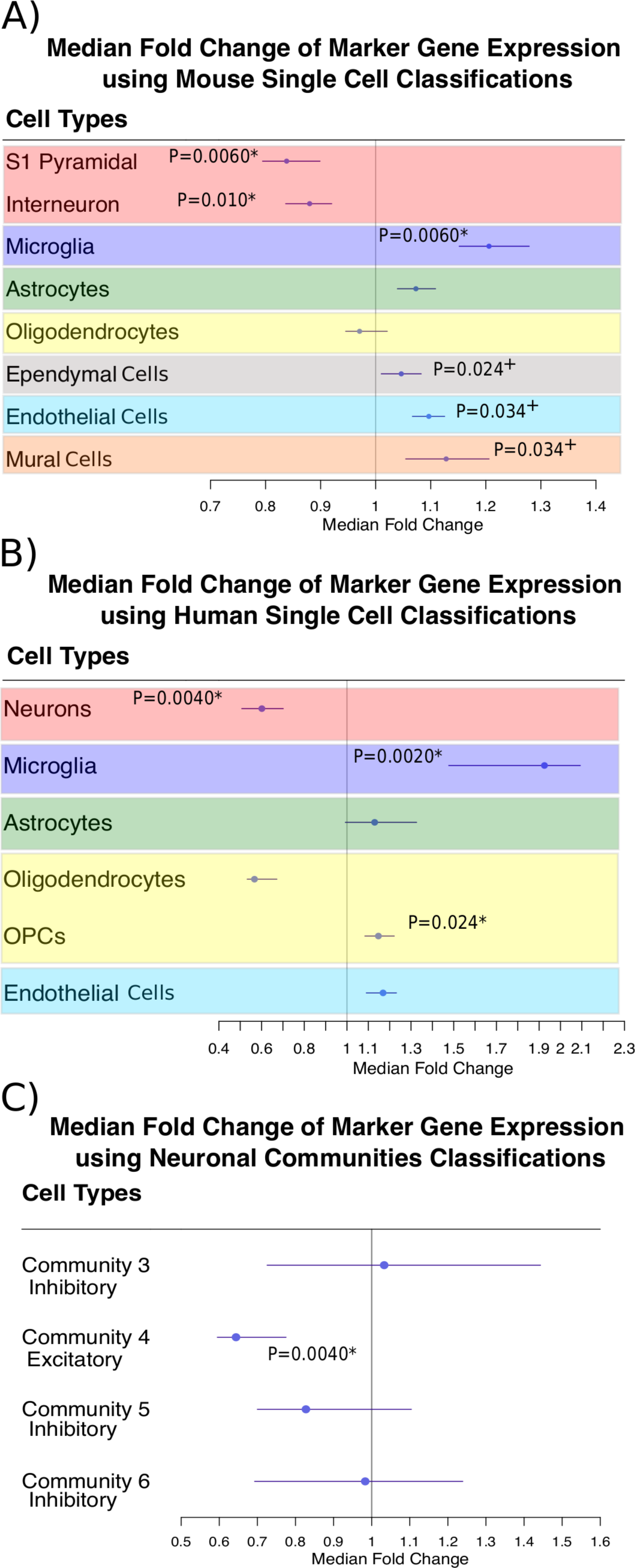
Plots of median expression fold change for cell type marker genes with bootstrapped 95% confidence intervals in the ASD versus control comparison. (A) For orthologous mouse marker genes. (B) For human marker genes. (C) For human neuronal community marker genes. Community 4 is an excitatory neuronal community and communities 3, 5, and 6 are inhibitory neuronal communities^24^. Significant p-values at an FDR<5% are marked by an asterisk (*) and marginally significant p-values at an FDR<5.1% are marked by a plus symbol (+). We also found consistent significant differences in analytical distributional tests but not in permutation-based distributional tests (data not shown). This could be due to the non-independence of expression of cell type markers (i.e. cell type gene markers will tend to be correlated in expression), ordinary interindividual variation in the relative abundance of these cell types, and the relatively underpowered nature of a 12 versus 12 permutation test.

Similar statistically significant results were observed, with even stronger effects, using the smaller set of cell type gene markers obtained from single cell transcriptomics in human cortex^24^ (Figure 2; Supplementary Table 3). Human cell type marker median gene expression for neurons was decreased in post-mortem human ASD cortex tissue by 30–49%. Conversely, microglia marker median expression was increased in ASD by 48–109°%, along with oligodendrocyte precursor cell (OPC) marker genes, whose median expression was increased in ASD by 8–22%. We then repeated this analysis for four previously defined excitatory or inhibitory neuronal subpopulations in human^24^. Gene markers of subpopulation 4, representing a type of excitatory neuron, were decreased in median expression in ASD brains by 22–40% (Figure 2C, Supplementary Table 4).

Having found that a majority of gene markers of several cortical cell types change in expression in ASD, we next asked whether these observations were more consistent with alterations in a specific pathway or cell state, or else with larger changes in cell type density. We would expect that alterations in specific pathways exclusively would produce expression changes amongst a minority subset of genes where expression of some markers (those contributing to the pathway or cell state) change while others do not. Instead, we found that expression of these marker genes shifted concordantly (Figure 3), suggesting cell type density changes in ASD.

**Figure 3.**
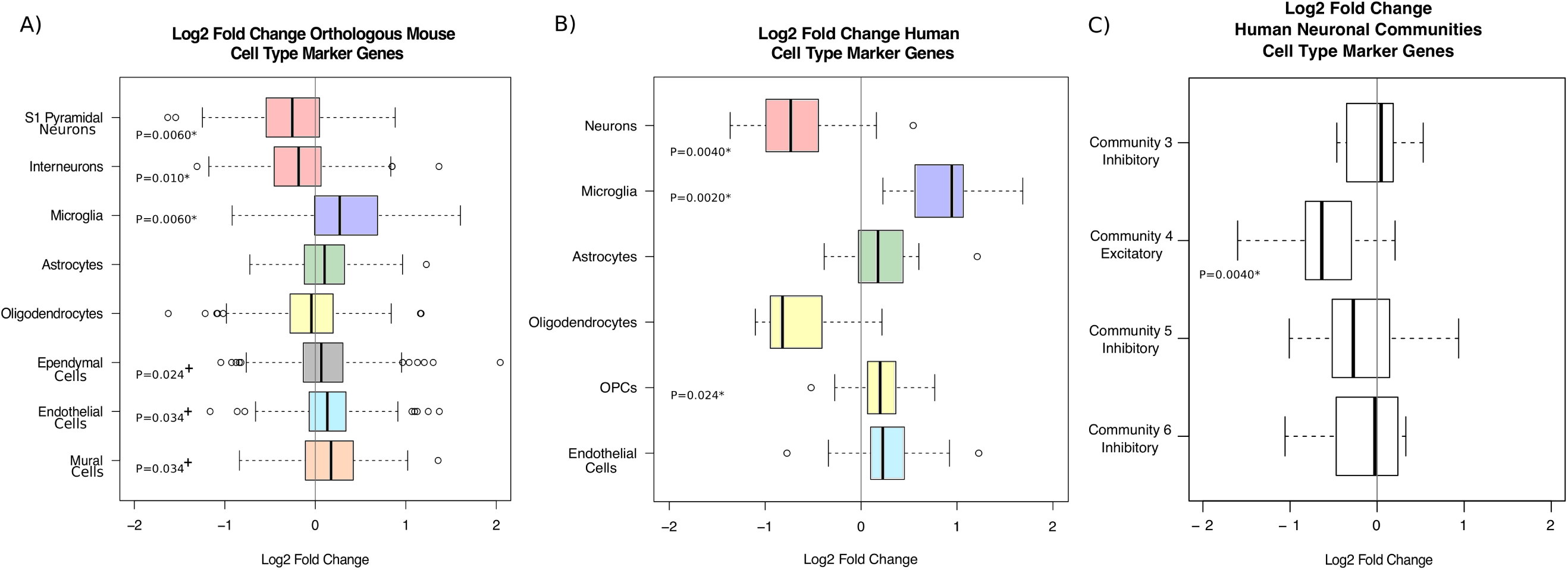
Boxplots representing the expression log_2_(fold change) distributions of cell type markers in ASD. (A) For orthologous mouse marker genes. (B) For human marker genes. (C) For human neuronal community marker genes. Community 4 is an excitatory neuronal community while communities 3, 5, and 6 are inhibitory neuronal communities^24^. Community 3, n=5 genes; community 4, n=23 genes; community 5, n=13 genes; community 6, n=8 genes. Interpreting the shapes of boxplots for sample sizes < 10 should be approached with caution^52^. The middle line represents the median, the left line is the 25th percentile (Q1), the right line is the 75th percentile (Q3), whiskers represent the minimum and maximum values excluding outliers, and outliers are depicted as individual points for values less than Q1–1.5*(Q3-Q1) and greater than Q3+1.5*(Q3-Q1). The asterisks (*) represent cell types significant at an FDR<5% and the plus symbols (+) represent those significant at an FDR<5.1%.

Expression changes of neuronal marker genes could reflect alterations not in cell number density, but in the density of subcellular compartments such as synapses. To consider this, we next examined whether protein products of neuronal marker genes that are localized in different neuronal compartments alter their mRNAs’ expression in ASD coordinately. S1 pyramidal and interneuron mouse markers were classified based on whether their protein products are localized in the nucleus of neurons or in dendrites and axons (Figure 4A, Supplementary Table 5). Mouse S1 pyramidal gene markers whose protein products are localized in the nucleus, and axons and dendrites, are similarly and significantly lower in transcript expression in ASD brains by 9–32°% and 6–32 %, respectively. The corresponding changes for mouse interneuron markers were 2–18 % and 6–24°%, respectively. Approximately three-quarters of genes for each of these localization categories show a decreased expression fold change (Figure 4B). These findings suggest that reduced expression of neuronal marker genes reflects decreased cell density.

**Figure 4.**
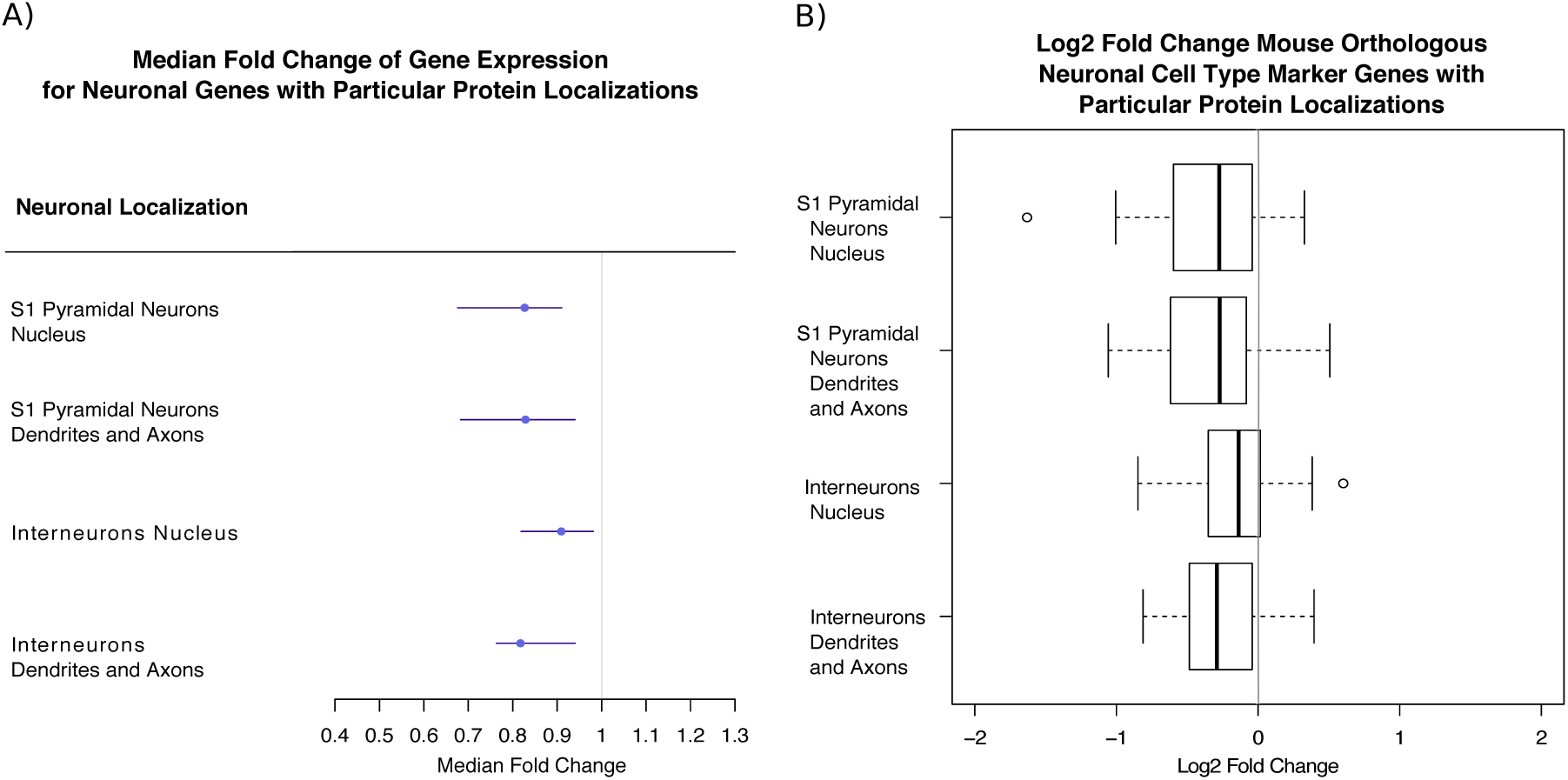
(A) Plots indicating the 95% confidence intervals of median fold change in ASD for orthologous mouse S1 pyramidal and interneuron marker genes with different protein product localizations. (B) Boxplot representing the distribution of expression log_2_(fold changes) in ASD of orthologous mouse S1 pyramidal and interneuron marker genes with different protein product localizations. All differences are significant at an FDR<5%.

## Discussion

In this study, the majority of neuronal and microglial cell marker genes exhibit coordinated expression level change in ASD brains. Previous enrichments of microglial markers in overexpressed-in-ASD ‘modules’ of co-expressed genes and enrichments of neuronal markers in underexpressed-in-ASD ‘modules’ of co-expressed genes^5^ were consistent with altered cellular processes besides changes in cell density. For example, increased microglial activation, observed in living young men with ASD^32^, might result in overexpression of M2 microglial activation genes even in the absence of a change in microglia density^33^. Furthermore, the enrichment of synaptic marker genes in under-expressed modules in ASD^5^ may be due to dysfunctional synaptic formation^9, 10^. Indeed, gene markers related to neurotransmission are underexpressed in adult ASD post-mortem cortex^34^. Nevertheless, that study failed to find differences in expression of neuronal and glial marker genes in ASD, probably due to the few markers used. Although there is some overlap between the cell type marker genes used here and microglia activation genes^33^ or neurotransmission genes^34^, we propose that the parsimonious interpretation of these results is that ASD brains contain alterations in cell density. We do so for the following reasons: (1) marker genes shift their expression together in the same direction in the ASD versus control comparison, rather than being driven by a minority subset as expected when only a specific pathway is altered, and (2) the median fold changes for transcripts encoding proteins localized in different neuronal compartments are all reduced, which is unexpected if only neuronal processes changed in their density.

For microglia, the issues of cell state and density may be correlated, because microglial activation can trigger proliferation^35, 36^. Microglial activation, increased density, and increased somal volume in ASD cortex have all been reported^37, 38^. The role of microglial activation and proliferation in ASD is unclear but increased microglia-neuronal spatial clustering in the dorsolateral prefrontal cortex in ASD suggests that active microglia are recruited to neurons^39^.

Decreases in neuronal density may not necessarily be caused by a reduction in relative cell number (i.e. the proportion of cell type nuclei versus all nuclei). We cannot exclude the possibility that these cell type marker gene expression changes in ASD may be due to differences in cell size, because gene expression of individual transcripts increases in proportion to cell volume^40, 41^. Hence, decreased neuronal density may be explained by less cytoplasmic space being taken up by neurons as a proportion of total cytoplasmic space.

Studies analyzing neuronal composition and density in ASD using immunohistochemistry and stereology have been conflicting, perhaps due to small sample sizes and the disorders’ heterogeneity. Patches of cortical disorganization and a small increase in neuronal density have been reported in children with ASD^16^. In the fusiform gyrus, reduced neuronal density, neuronal numbers, and cell volumes were reported in ASD, but these changes were not detected in whole cortical grey matter or primary visual cortex^42^. In the dorsolateral and mesial prefrontal cortex of children, an increase in neuronal counts but not in neuronal volume was observed^43^. However, in Brodmann areas 44 and 45, smaller pyramidal neurons, albeit similar pyramidal neuronal numbers, were reported in ASD^44^. An interesting possibility is that the underlying structure of neurons in the cortex is different in ASD, resulting in decreased density in the cortical section sequenced. Increased center-to-center minicolumn width has been observed in different cortical areas in ASD, including a trend for increased width in the primary auditory cortex (Brodmann 41, overlapping the area used in this study)^19^. Wider minicolumn spacing in ASD may explain decreased expression of neuronal markers, as it would imply a lower density of neurons.

These results are consistent with previous transcriptomic studies, and expand on them. These have shown an enrichment of interneuron and glutamatergic neuron markers in an ASD underexpressed gene module in the cortex and enrichment of activated microglia and astrocyte markers in an ASD overexpressed module^5, 7^. Later work reported an over-representation of overexpressed genes in immune cells and astrocytes and an over-representation of underexpressed genes in interneurons and putative cortical projection neurons in the Voineagu et al. dataset^8^. Another study also observed overexpression of microglia and astrocyte markers and underexpression of neuronal markers in the prefrontal cortex, but used few markers^45^. Here, we used objectively defined gene markers, allowing us to pinpoint excitatory neuronal markers as being specifically downregulated. We also included non-neuronal and glial cell types, which previous studies did not include. Instead of analyzing the enrichment of cell type markers in underexpressed and overexpressed genes in ASD, we tested whether these markers change expression as a group. For the first time, to our knowledge, this method allows us make inferences about the composition of cell types in the ASD brain.

The transcriptomic data used in this study was obtained from the superior temporal gyrus. Since this brain region is important for auditory processing and speech^46–48^ and has been implicated in ASD^49, 50^, it is important to determine how and if cell type density changes here lead to behavioral deficits. Although these gene expression patterns may be regionally specific, there is strong overlap between differentially expressed genes in the temporal and frontal cortex, suggesting that gene expression changes in the cortex might be global and not representative of a particular region^5^.

In this study, we used a cell type transcriptomic approach to infer cellular composition differences in ASD cortex. The median gene expression of inhibitory or excitatory neuronal marker genes was significantly reduced in ASD. This was also the case when subdividing mouse orthologous neuronal markers based on the subcellular localization of the proteins they encode. The median gene expression of microglia markers was significantly increased in ASD. These gene expression patterns suggest changes in cell type number and/or volume density. Given the conflicting results in published immunohistochemistry and stereology studies, understanding the underlying causes of these global gene expression signatures in ASD should be prioritized. Groups with expertise in stereology might now independently validate this inference from markers to cell type density. Since the patients used in this analysis were a subset displaying strong characteristic differential gene expression changes in ASD, future integrated transcriptomics and stereological studies could compare density changes in post-mortem tissue with and without these distinctive gene expression changes. Moreover, this study provides a proof of concept for using single cell transcriptomic analyses derived marker genes to study cell type composition changes in disease, and could be easily extended to the study of other diseases such as schizophrenia, where cell type marker enrichment has been reported in gene co-expression modules^51^.

## Acknowledgements

RBM was supported by the Clarendon Fund; RBM, TGB, and CPP were supported by the Medical Research Council; and TGB was supported by the Human Brain Project. The research leading to these results has received funding from the European Union Seventh Framework Programme (FP7/2007–2013) under grant agreement no. 604102 (Human Brain Project).Conflict of Interest

The authors declare no financial conflicts of interest.

## Supplementary information for

**Supplementary Table 1.** Cell Types and Number of Genes per Cell Type for Mouse Orthologous and Human Marker Genes

| Cell Type (from mouse single cell transcriptomics) | Number of Genes in Publication (Zeisel et al.) | Number of Genes in Analysis |
| --- | --- | --- |
| **S1 Pyramidal Neurons** | 294 | 180 |
| **Interneurons** | 365 | 264 |
| **Microglia** | 436 | 234 |
| **Astrocytes** | 240 | 185 |
| **Oligodendrocytes** | 453 | 348 |
| **Ependymal Cells** | 484 | 266 |
| **Endothelial Cells** | 353 | 237 |
| **Mural Cells** | 155 | 95 |
| Cell Type (from human single cell transcriptomics) | Number of Genes in Publication (Darmanis et al.) | Number of Genes in Analysis |
| **Neurons** | 21 | 21 |
| **Microglia** | 21 | 13 |
| **Astrocytes** | 21 | 19 |
| **Oligodendrocytes** | 21 | 19 |
| **Oligodendrocyte Precursor Cells (OPCs)** | 21 | 15 |
| **Endothelial** | 21 | 16 |

**Supplementary Table 2.** Median expression log_2_(FC) of Mouse Orthologous Marker Genes

| Cell Type | Number of Genes | Median log<sub>2</sub>(FC) | Median FC | Empirical p-value | log<sub>2</sub>(CI) 2.5% | log<sub>2</sub>(CI) 97.5% | CI 2.5% | CI 97.5% |
| --- | --- | --- | --- | --- | --- | --- | --- | --- |
| **S1 Pyramidal Neurons** | 180 | -0.25 | 0.84 | 0.0060 * | -0.33 | -0.15 | 0.79 | 0.90 |
| **Interneurons** | 264 | -0.18 | 0.88 | 0.010 * | -0.26 | -0.12 | 0.84 | 0.92 |
| **Microglia** | 234 | 0.27 | 1.21 | 0.0060 * | 0.20 | 0.36 | 1.15 | 1.28 |
| **Astrocytes** | 185 | 0.10 | 1.07 | 0.058 | 0.06 | 0.15 | 1.04 | 1.11 |
| **Oligodendrocytes** | 348 | -0.04 | 0.97 | 0.72 | -0.08 | 0.03 | 0.95 | 1.02 |
| **Ependymal Cells** | 266 | 0.07 | 1.05 | 0.024 <sup>+</sup> | 0.01 | 0.12 | 1.01 | 1.08 |
| **Endothelial Cells** | 237 | 0.13 | 1.10 | 0.034 <sup>+</sup> | 0.09 | 0.17 | 1.07 | 1.13 |
| **Mural Cells** | 95 | 0.17 | 1.13 | 0.034 <sup>+</sup> | 0.08 | 0.27 | 1.05 | 1.21 |

**Supplementary Table 3.** Median expression log_2_(FC) of Human Marker Genes

| Cell Type | Number of Genes | Median $\log_2(\text{FC})$ | Median FC | Empirical p-value | $\log_2(\text{CI})$ 2.5% | $\log_2(\text{CI})$ 97.5% | CI 2.5% | CI 97.5% |
| --- | --- | --- | --- | --- | --- | --- | --- | --- |
| Neurons | 21 | -0.73 | 0.60 | 0.0040 * | -0.98 | -0.51 | 0.51 | 0.70 |
| Microglia | 13 | 0.95 | 1.93 | 0.0020 * | 0.56 | 1.07 | 1.48 | 2.09 |
| Astrocytes | 19 | 0.18 | 1.13 | 0.40 | -0.01 | 0.41 | 0.99 | 1.33 |
| Oligodendrocytes | 19 | -0.82 | 0.57 | 0.064 | -0.91 | -0.57 | 0.53 | 0.67 |
| OPCs | 15 | 0.20 | 1.15 | 0.024 * | 0.12 | 0.29 | 1.08 | 1.22 |
| Endothelial Cells | 16 | 0.23 | 1.17 | 0.30 | 0.13 | 0.30 | 1.09 | 1.23 |

**Supplementary Table 4.** Median expression log_2_(FC) of Human Neuronal Communities Marker Genes

| Neuronal Community | Number of Genes | Median $\log_2(\text{FC})$ | Median FC | Empirical p-value | $\log_2(\text{CI})$ 2.5% | $\log_2(\text{CI})$ 97.5% | CI 2.5% | CI 97.5% |
| --- | --- | --- | --- | --- | --- | --- | --- | --- |
| 3 | 5 | 0.05 | 1.03 | 0.82 | -0.46 | 0.53 | 0.73 | 1.44 |
| 4 | 23 | -0.64 | 0.64 | 0.0040 * | -0.75 | -0.37 | 0.60 | 0.78 |
| 5 | 13 | -0.27 | 0.83 | 0.056 | -0.52 | 0.14 | 0.70 | 1.10 |
| 6 | 8 | -0.02 | 0.98 | 0.86 | -0.53 | 0.31 | 0.69 | 1.24 |

**Supplementary Table 5.**
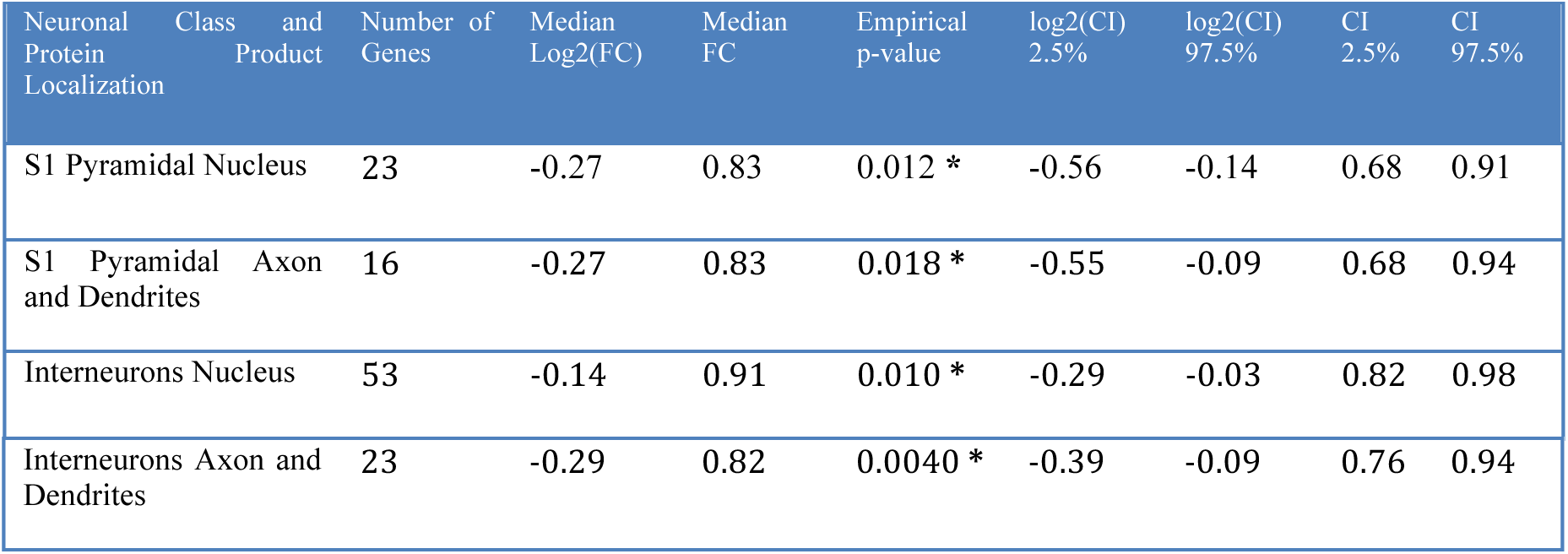
Median expression log_2_(FC) of Mouse Orthologous Neuronal Marker Genes Subdivided by Protein Product Localization

## References

1. American Psychiatric Association. Diagnostic and statistical manual of mental disorders: DSM-5. Fifth edition. edn. American Psychiatric Publishing: Washington, DC, 2013, xliv, 947 pagespp.

2. Hallmayer J, Cleveland S, Torres A, Phillips J, Cohen B, Torigoe T et al. Genetic heritability and shared environmental factors among twin pairs with autism. Arch Gen Psychiatry 2011; 68(11): 1095–1102.

3. Rosenberg RE, Law JK, Yenokyan G, McGready J, Kaufmann WE, Law PA. Characteristics and concordance of autism spectrum disorders among 277 twin pairs. Arch Pediatr Adolesc Med 2009; 163(10): 907–914.

4. Ronemus M, Iossifov I, Levy D, Wigler M. The role of de novo mutations in the genetics of autism spectrum disorders. Nat Rev Genet 2014; 15(2): 133–141.

5. Voineagu I, Wang X, Johnston P, Lowe JK, Tian Y, Horvath S et al. Transcriptomic analysis of autistic brain reveals convergent molecular pathology. Nature 2011; 474(7351): 380–384.

6. Irimia M, Weatheritt RJ, Ellis JD, Parikshak NN, Gonatopoulos-Pournatzis T, Babor M et al. A highly conserved program of neuronal microexons is misregulated in autistic brains. Cell 2014; 159(7): 1511–1523.

7. Parikshak NN, Luo R, Zhang A, Won H, Lowe JK, Chandran V et al. Integrative functional genomic analyses implicate specific molecular pathways and circuits in autism. Cell 2013; 155(5): 1008–1021.

8. Xu X, Wells AB, O’Brien DR, Nehorai A, Dougherty JD. Cell type-specific expression analysis to identify putative cellular mechanisms for neurogenetic disorders. The Journal of neuroscience: the official journal of the Society for Neuroscience 2014; 34(4): 1420–1431.

9. Rubenstein JL, Merzenich MM. Model of autism: increased ratio of excitation/inhibition in key neural systems. Genes Brain Behav 2003; 2(5): 255–267.

10. Jamain S, Quach H, Betancur C, Rastam M, Colineaux C, Gillberg IC et al. Mutations of the X-linked genes encoding neuroligins NLGN3 and NLGN4 are associated with autism. Nat Genet 2003; 34(1): 27–29.

11. Kreczmanski P, Heinsen H, Mantua V, Woltersdorf F, Masson T, Ulfig N et al. Volume, neuron density and total neuron number in five subcortical regions in schizophrenia. Brain 2007; 130(Pt 3): 678–692.

12. Ribeiro AA, Davis C, Gabella G. Estimate of size and total number of neurons in superior cervical ganglion of rat, capybara and horse. Anat Embryol (Berl) 2004; 208(5): 367–380.

13. Ramos R, Requena V, Diaz F, Villena A, Perez de Vargas I. Evolution of neuronal density in the ageing thalamic reticular nucleus. Mech Ageing Dev 1995; 83(1): 21–29.

14. Casanova MF, Buxhoeveden DP, Switala AE, Roy E. Neuronal density and architecture (Gray Level Index) in the brains of autistic patients. J Child Neurol 2002; 17(7): 515–521.

15. Lawrence YA, Kemper TL, Bauman ML, Blatt GJ. Parvalbumin-, calbindin-, and calretinin-immunoreactive hippocampal interneuron density in autism. Acta Neurol Scand 2010; 121(2): 99–108.

16. Stoner R, Chow ML, Boyle MP, Sunkin SM, Mouton PR, Roy S et al. Patches of disorganization in the neocortex of children with autism. N Engl J Med 2014; 370(13): 1209–1219.

17. Bailey A, Luthert P, Dean A, Harding B, Janota I, Montgomery M et al. A clinicopathological study of autism. Brain 1998; 121 (Pt 5): 889–905.

18. Casanova MF, Buxhoeveden DP, Switala AE, Roy E. Minicolumnar pathology in autism. Neurology 2002; 58(3): 428–432.

19. McKavanagh R, Buckley E, Chance SA. Wider minicolumns in autism: a neural basis for altered processing? Brain 2015; 138(Pt 7): 2034–2045.

20. Doyle JP, Dougherty JD, Heiman M, Schmidt EF, Stevens TR, Ma G et al. Application of a translational profiling approach for the comparative analysis of CNS cell types. Cell 2008; 135(4): 749–762.

21. Sandberg R. Entering the era of single-cell transcriptomics in biology and medicine. Nature methods 2014; 11(1): 22–24.

22. Usoskin D, Furlan A, Islam S, Abdo H, Lonnerberg P, Lou D et al. Unbiased classification of sensory neuron types by large-scale single-cell RNA sequencing. Nature neuroscience 2015; 18(1): 145–153.

23. Zeisel A, Munoz-Manchado AB, Codeluppi S, Lonnerberg P, La Manno G, Jureus A et al. Brain structure. Cell types in the mouse cortex and hippocampus revealed by single-cell RNA-seq. Science 2015; 347(6226): 1138–1142.

24. Darmanis S, Sloan SA, Zhang Y, Enge M, Caneda C, Shuer LM et al. A survey of human brain transcriptome diversity at the single cell level. Proceedings of the National Academy of Sciences of the United States of America 2015; 112(23): 7285–7290.

25. UniProt C. UniProt: a hub for protein information. Nucleic Acids Res 2015; 43(Database issue): D204–212.

26. Roncaglia P, Martone ME, Hill DP, Berardini TZ, Foulger RE, Imam FT et al. The Gene Ontology (GO) Cellular Component Ontology: integration with SAO (Subcellular Anatomy Ontology) and other recent developments. J Biomed Semantics 2013; 4(1): 20.

27. Martin KC, Zukin RS. RNA trafficking and local protein synthesis in dendrites: an overview. The Journal of neuroscience: the official journal of the Society for Neuroscience 2006; 26(27): 7131–7134.

28. Willis DE, Twiss JL. Regulation of protein levels in subcellular domains through mRNA transport and localized translation. Mol Cell Proteomics 2010; 9(5): 952–962.

29. Kinsella RJ, Kahari A, Haider S, Zamora J, Proctor G, Spudich G et al. Ensembl BioMarts: a hub for data retrieval across taxonomic space. Database (Oxford) 2011; 2011: bar030.

30. R Core Team. R: A language and environment for statistical computing. R Foundation for Statistical Computing2014.

31. Benjamini Y, Hochberg Y. Controlling the False Discovery Rate: A Practical and Powerful Approach to Multiple Testing. Journal of the Royal Statistical Society Series B (Methodological) 1995; 57(1): 289–300.

32. Suzuki K, Sugihara G, Ouchi Y, Nakamura K, Futatsubashi M, Takebayashi K et al. Microglial activation in young adults with autism spectrum disorder. JAMA Psychiatry 2013; 70(1): 49–58.

33. Gupta S, Ellis SE, Ashar FN, Moes A, Bader JS, Zhan J et al. Transcriptome analysis reveals dysregulation of innate immune response genes and neuronal activity-dependent genes in autism. Nature communications 2014; 5: 5748.

34. van de Lagemaat LN, Nijhof B, Bosch DG, Kohansal-Nodehi M, Keerthikumar S, Heimel JA. Age-related decreased inhibitory vs. excitatory gene expression in the adult autistic brain. Front Neurosci 2014; 8: 394.

35. Kettenmann H, Hanisch UK, Noda M, Verkhratsky A. Physiology of microglia. Physiol Rev 2011; 91(2): 461–553.

36. Graeber MB, Tetzlaff W, Streit WJ, Kreutzberg GW. Microglial cells but not astrocytes undergo mitosis following rat facial nerve axotomy. Neuroscience letters 1988; 85(3): 317–321.

37. Morgan JT, Chana G, Pardo CA, Achim C, Semendeferi K, Buckwalter J et al. Microglial activation and increased microglial density observed in the dorsolateral prefrontal cortex in autism. Biol Psychiatry 2010; 68(4): 368–376.

38. Tetreault NA, Hakeem AY, Jiang S, Williams BA, Allman E, Wold BJ et al. Microglia in the cerebral cortex in autism. J Autism Dev Disord 2012; 42(12): 2569–2584.

39. Morgan JT, Chana G, Abramson I, Semendeferi K, Courchesne E, Everall IP. Abnormal microglial-neuronal spatial organization in the dorsolateral prefrontal cortex in autism. Brain Res 2012; 1456: 72–81.

40. Wu CY, Rolfe PA, Gifford DK, Fink GR. Control of transcription by cell size. PLoS Biol 2010; 8(11): e1000523.

41. Zhurinsky J, Leonhard K, Watt S, Marguerat S, Bahler J, Nurse P. A coordinated global control over cellular transcription. Curr Biol 2010; 20(22): 2010–2015.

42. van Kooten IA, Palmen SJ, von Cappeln P, Steinbusch HW, Korr H, Heinsen H et al. Neurons in the fusiform gyrus are fewer and smaller in autism. Brain 2008; 131(Pt 4): 987–999.

43. Courchesne E, Mouton PR, Calhoun ME, Semendeferi K, Ahrens-Barbeau C, Hallet MJ et al. Neuron number and size in prefrontal cortex of children with autism. JAMA 2011; 306(18): 2001–2010.

44. Jacot-Descombes S, Uppal N, Wicinski B, Santos M, Schmeidler J, Giannakopoulos P et al. Decreased pyramidal neuron size in Brodmann areas 44 and 45 in patients with autism. Acta Neuropathol 2012; 124(1): 67–79.

45. Edmonson C, Ziats MN, Rennert OM. Altered glial marker expression in autistic post-mortem prefrontal cortex and cerebellum. Mol Autism 2014; 5(1): 3.

46. Hillis AE, Wityk RJ, Tuffiash E, Beauchamp NJ, Jacobs MA, Barker PB et al. Hypoperfusion of Wernicke’s area predicts severity of semantic deficit in acute stroke. Ann Neurol 2001; 50(5): 561–566.

47. Mesgarani N, Cheung C, Johnson K, Chang EF. Phonetic feature encoding in human superior temporal gyrus. Science 2014; 343(6174): 1006–1010.

48. Pickles JO. An introduction to the physiology of hearing. 4th edn. Emerald: Bingley, 2012, xxiii, 430 p., 434 p. of platespp.

49. Gendry Meresse I, Zilbovicius M, Boddaert N, Robel L, Philippe A, Sfaello I et al. Autism severity and temporal lobe functional abnormalities. Ann Neurol 2005; 58(3): 466–469.

50. Bigler ED, Mortensen S, Neeley ES, Ozonoff S, Krasny L, Johnson M et al. Superior temporal gyrus, language function, and autism. Dev Neuropsychol 2007; 31(2): 217–238.

51. Torkamani A, Dean B, Schork NJ, Thomas EA. Coexpression network analysis of neural tissue reveals perturbations in developmental processes in schizophrenia. Genome Res 2010; 20(4): 403–412.

52. Krzywinski M, Altman N. Visualizing samples with box plots. Nature methods 2014; 11(2): 119–120.

